# Human Hippocampal Theta-Gamma Phase-Amplitude Coupling Signals the Distance to Remembered Goals

**DOI:** 10.64898/2026.10.08.757608

**Authors:** Zimo Huang, Paris Goumas, James A Bisby, Roman Rodionov, Beate Diehl, Andrew W McEvoy, Matthew C Walker, Umesh Vivekananda, Neil Burgess, Daniel Bush

## Abstract

In mammals, flexible navigation is believed to rely on a cognitive map of known locations supported by the hippocampus and surrounding medial temporal lobe structures. Hippocampal place cells typically exhibit spatial firing fields within a given environment. Moreover, their firing is modulated by the movement-related theta rhythm such that ‘theta sweeps’ of place cell activity encode the trajectory towards a hidden reward location during goal-directed navigation. Here, we attempted to detect signatures of theta sweeps in the human hippocampal local field potential (LFP) recorded with intracranial EEG. Specifically, we hypothesised that sequences of locations along a planned trajectory encoded by distinct cell assemblies are associated with gamma activity at successive theta phases. To test this hypothesis, we asked N=9 patients with epilepsy to perform a spatial navigation task in desktop virtual reality while we recorded LFP signals from depth electrode contacts implanted across the medial temporal lobe. Consistent with our predictions, we found an increase in hippocampal theta-gamma phase-amplitude coupling, a decrease in gamma power, and theta phase precession of gamma activity as hidden goal locations were approached and the sequence of locations remaining to be traversed became shorter. This suggests that goal-directed hippocampal theta sweeps during navigation are reflected in LFP theta-gamma phase-amplitude coupling, and that these neural mechanisms of model-based planning are conserved across species.

## Introduction

Human behaviour is remarkable in its capacity for flexibility and generalisation, extrapolating from limited experience to rapidly optimise performance on novel tasks. This is thought to reflect the construction of an internal model of the world, or “cognitive map”, which can be used to evaluate the outcome of specific sequences of behaviour (Tolman, 1948; O’Keefe and Nadel, 1978; Burgess et al., 2002; Daw et al., 2005; Behrens et al., 2018). One canonical example comes from spatial navigation, for which the underlying neuronal representations are found in the hippocampus and surrounding medial temporal lobe (Moser et al., 2017). Most notably, place cells in the hippocampus proper are typically active in single, restricted regions of space known as the corresponding ‘place field’ (O’Keefe and Dostrovsky, 1971). As such, hippocampal population activity encodes an animal’s current location within any given environment (Brown et al., 1998) across a range of spatial scales, with smaller place fields in the dorsal (posterior) hippocampus and larger place fields in the ventral (anterior) hippocampus (Kjelstrup et al., 2008; Strange et al., 2014). In addition, place cell activity is strongly modulated by the 6-10Hz theta rhythm during movement (Fox et al., 1986) and exhibits theta phase precession: firing at progressively earlier theta phases as their place fields are traversed (O’Keefe and Recce, 1993). At the population level, this creates ‘theta sweeps’ of place cell activity within each oscillatory cycle that typically encode a sequence of locations along the current movement trajectory (Burgess et al., 1994; Skaggs et al., 1996; Johnson and Redish, 2007; Vollan et al., 2025). More recently, it has been demonstrated that theta sweeps are oriented towards hidden reward locations during goal-directed navigation (Tang et al., 2026; Yu et al., 2026), and that the length of theta sweep trajectories correlates with distance to the current goal location (Wikenheiser and Redish, 2015; Yu et al., 2026). This suggests that theta sweeps reflect a mechanism by which the hippocampal cognitive map is used to support flexible navigation.

Place cell firing patterns (Ekstrom et al., 2003), theta phase precession (Qasim et al., 2021), and theta phase specific firing during sequential learning tasks (Reddy et al., 2021) have also been reported in single unit recordings from the human hippocampus. Identifying theta sweeps, however, requires large numbers of simultaneously recorded neurons for decoding, which is difficult to achieve in humans. Instead, we hypothesised that it might be possible to identify signatures of goal-directed theta sweeps in the hippocampal local field potential (LFP). Specifically, we propose that sequences of locations along a planned trajectory are encoded by distinct cell assemblies associated with gamma activity at successive theta phases (Senior et al., 2008), consistent with models of working memory function (Lisman and Idiart, 1995; Jensen and Lisman, 1998; Lisman, 2005). As the goal location is approached, and the sequence of locations remaining to be traversed becomes shorter, this predicts an increase in phase-amplitude coupling as gamma activity becomes more narrowly distributed in the theta cycle (Heusser et al., 2016). Previously, we verified this hypothesis using non-invasive MEG recordings from healthy participants completing an abstract goal-directed navigation paradigm (Huang et al., 2026). Here, we sought to replicate and extend these findings by examining direct intracranial EEG recordings from the hippocampus in pre-surgical epilepsy patients completing a more ethologically valid spatial navigation task (Doeller et al., 2008; Bush et al., 2017; Vivekananda et al., 2021). Specifically, investigated the relationship between hippocampal theta-gamma phase-amplitude coupling, gamma power, and the preferred theta phase of gamma activity during navigation to both visible and hidden goal locations in a desktop virtual reality (VR) environment.

## Results

### Behavioural Data

We analysed intracranial EEG recordings from depth electrodes targeting the medial temporal lobe of nine epilepsy patients completing a spatial memory task in desktop VR (Figure 1; Bush et al., 2017; Vivekananda et al., 2021; see Supplementary Table 1 for patient demographics). In this task, participants are first asked to memorise the location of four visible objects in a square, featureless environment across several ‘encoding’ trials in which only one object is visible at a time (see Methods for further details). Next, they are shown an image of one object, then placed back at a random location and orientation in the environment and asked to navigate to the remembered location of that object and make a response. This provides a continuous measurement of memory accuracy, as the distance ‘error’ between the response location and true location of the corresponding object. An examination of behavioural performance indicated that patients could accurately encode object locations and use those to support memory guided navigation. Specifically, the average distance error across patients was Δ*d* = 31.0 ± 15.4 virtual metres (mean ± SD, range of 10.5-51.1 vm), equivalent to a normalised performance of 0.725 ± 0.166 (see Methods for further details) and significantly better than chance (∼51.9 vm, t(8) = 4.05, *p* < 0.005).

**Figure 1:**
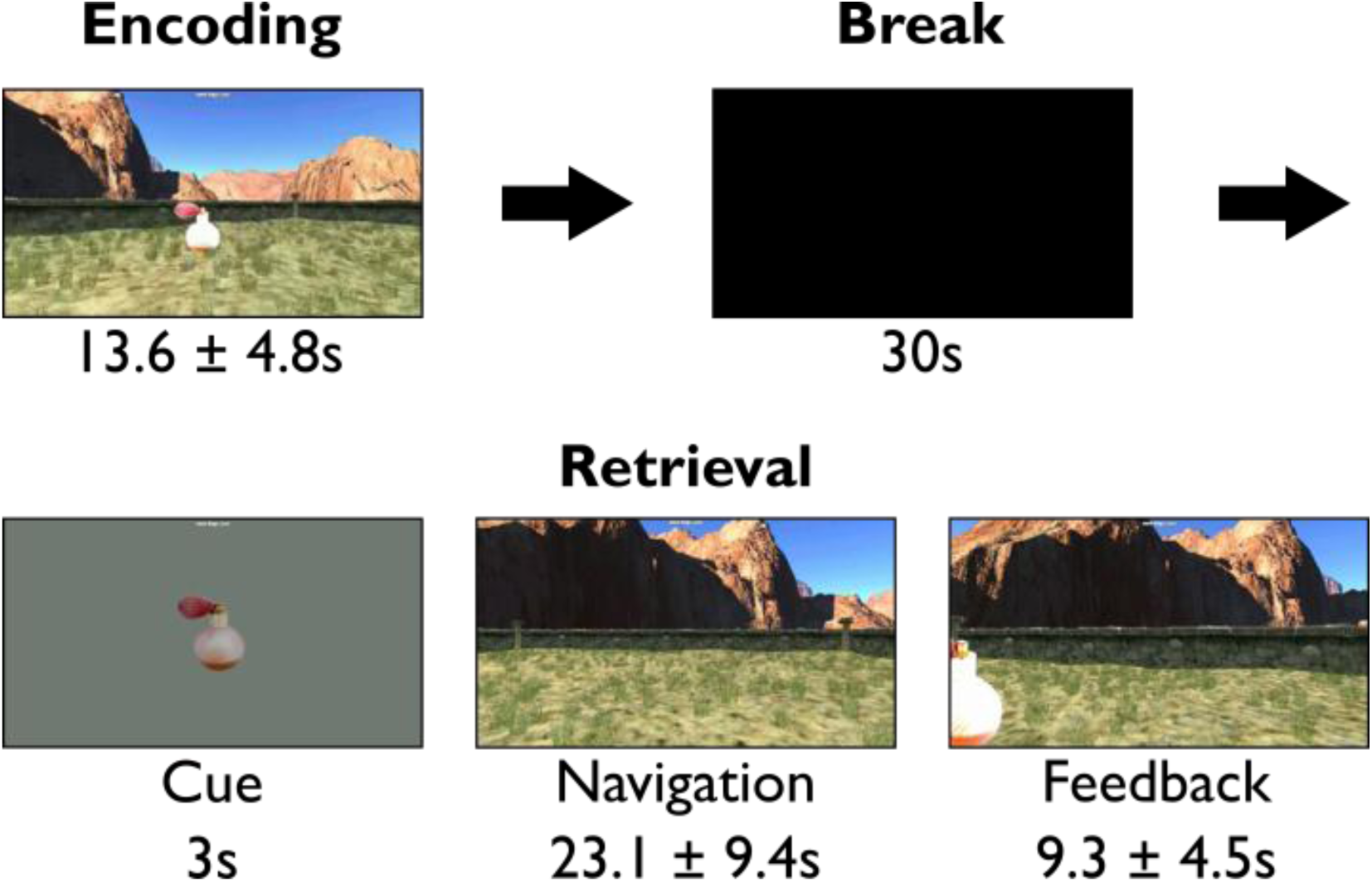
Spatial memory task in desktop virtual reality (VR; Doeller et al., 2008; Bush et al., 2017). Throughout the task, participants navigated freely in a square environment defined by a uniform floor texture and surrounding boundary with extra-maze cues projected at infinity for orientation. In each of twenty encoding trials (five trials for each of four distinct objects presented in a pseudorandom order), participants were asked to learn the location of a visible object and then collide with the visible object to proceed to the next trial. After a short break, their memory for those object locations was tested across twenty retrieval trials (five trials for each object presented in a pseudorandom order). Each retrieval trial began with a cue period in which an image of one object was presented, before participants were placed at a random location and orientation within the environment and asked to navigate to the remembered location of that object and make a response. Once they had done so, the object appeared in its correct location to provide feedback on their performance, and they were asked to collide with the (now visible) object to proceed to the next trial. Inset timings indicate the mean ± SD duration of each task phase across participants. Adapted from Bush et al., (2017).

Building on our previous MEG findings (Huang et al., 2026) and observations of goal-directed theta sweeps in rodents (Wikenheiser et al., 2015; Tang et al., 2026; Yu et al., 2026), we hypothesised that navigation in this task would be supported by hippocampal theta-gamma phase-amplitude coupling. According to this coding scheme, the sequence of locations that need to be traversed to reach the goal is encoded by distinct cell assemblies associated with gamma activity at successive theta phases. As the goal is approached during movement, this predicts that theta-gamma phase-amplitude coupling will increase, gamma power will decrease, and – if the initial theta phase of gamma activity remains fixed - the preferred theta phase of gamma activity will shift progressively earlier, as the remaining sequence of locations becomes shorter (Figure 2). Moreover, each of these effects should be more pronounced in the posterior hippocampus, where place fields are typically smaller and the spatial representation therefore changes more rapidly during movement (Kjelstrup et al., 2008; Liu et al., 2023).

**Figure 2:**
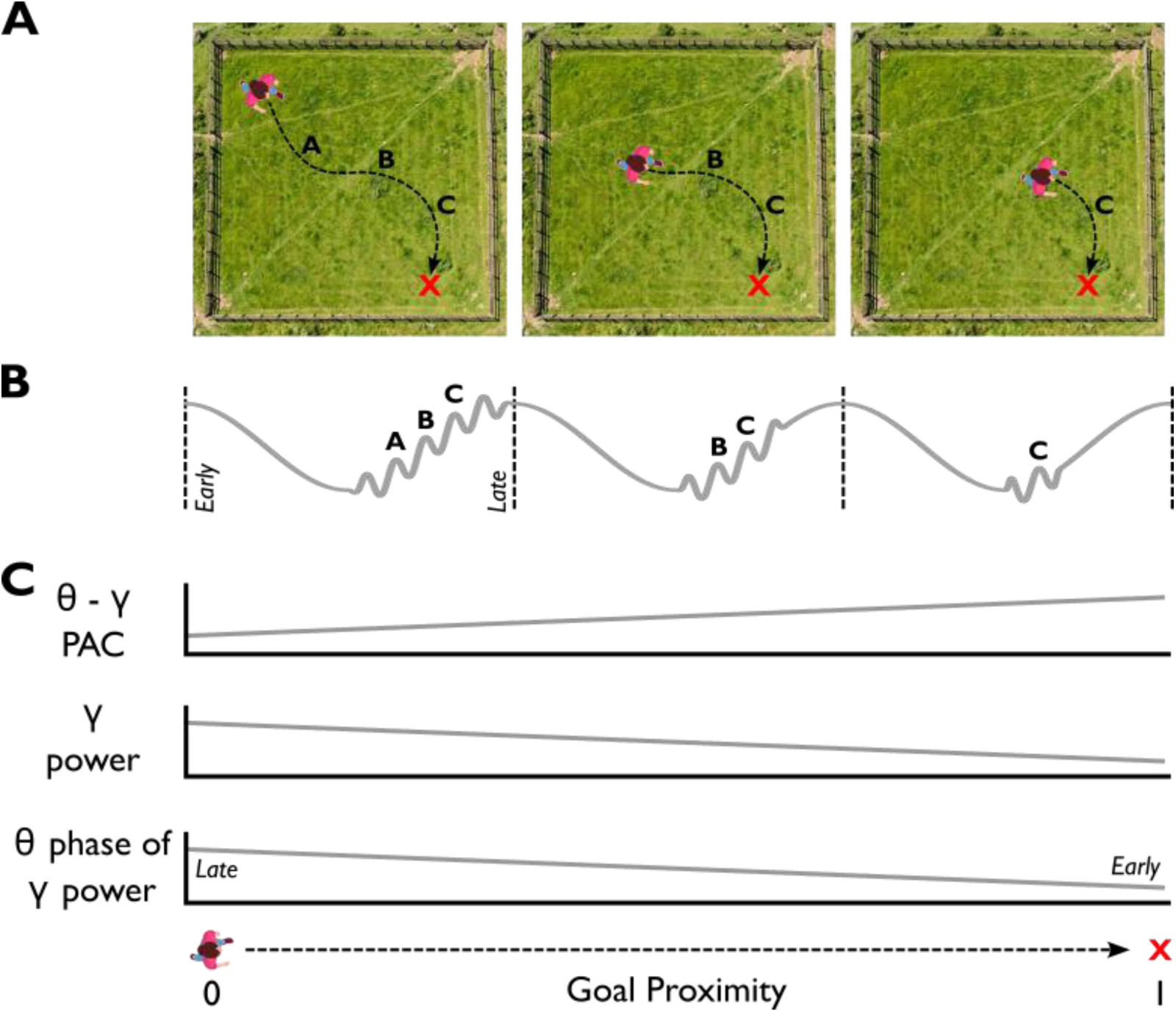
Experimental predictions. **(A)** The planned route to a goal location (marked with a red cross) passes through a sequence of locations (A, B, C) that are encoded by distinct hippocampal place cell assemblies. **(B)** If hippocampal theta sequences represent this intended route (Wikenheiser et al., 2015; Tang et al., 2026; Yu et al., 2026), then those place cell assemblies - which are associated with gamma activity in the local field potential (black line) - will be active at successive phases of each theta cycle. If hippocampal theta sequences terminate at the goal location, then the duration of gamma activity in each theta cycle will become progressively shorter as the goal is approached. **(C)** As such, increasing proximity to the goal will be associated with: (i) an increase in theta-gamma phase-amplitude coupling (θ – γ PAC), as the distribution of gamma activity across theta phases becomes narrower (top panel); (ii) a decrease in gamma power, as the duration of gamma activity in each theta cycle becomes shorter (middle panel); and (iii) the preferred theta phase of gamma power becoming progressively earlier, reflecting theta phase precession of the underlying place cell population s (bottom panel).

To test this hypothesis, we first identified periods of continuous translational movement that lasted >1s from throughout the task. Next, we divided those movement epochs into non-overlapping 1s windows and computed normalised proximity to the visible or remembered goal location (during encoding and retrieval trials, respectively) for each window. On average, patients completed a total of 119 ± 55 (range of 47-204) of these movement epochs across the whole task (with a total of 170 ± 68 non-overlapping 1s sliding windows, range of 63-258), with 101 ± 51.6 (range of 42-184) artefact-free epochs (144 ± 68 non-overlapping sliding windows, range 49-232) included in subsequent analyses (see Supplementary Table 2 for further details).

### Hippocampal Theta-Gamma Phase-Amplitude Coupling Increases with Proximity to the Goal during Memory-guided Navigation

First, we set out to examine whether dynamic changes in cross-frequency coupling between theta phase and gamma amplitude covary with distance from the goal during navigation. To do so, we conducted a linear regression between normalised goal distance and estimates of theta-gamma phase-amplitude coupling (TG-PAC) across hippocampal electrode contacts in non-overlapping 1 s time windows throughout movement (see Methods for further details). Initially, we focussed on the 2-5 Hz ‘low theta’ and 70-140 Hz ‘high gamma’ bands described by previous studies (Bush et al., 2017; Daume et al., 2024; Huang et al., 2026). Across both encoding and retrieval, we found that TG-PAC increased with proximity to the (visible and remembered) goal location (t(8) = 4.84, *p* = 0.00129, one-sample t-test). Subsequent analysis indicated that this effect was primarily driven by memory-guided navigation during retrieval (t(8) = 3.80, *p* = 0.005, one-sample t-test), but not visually guided navigation during encoding (t(8) = 1.05, *p* = 0.321), although the difference between encoding and retrieval trials was not significant (t(8) = -1.86, *p* = 0.1, paired t-test; Figure 3A).

**Figure 3:**
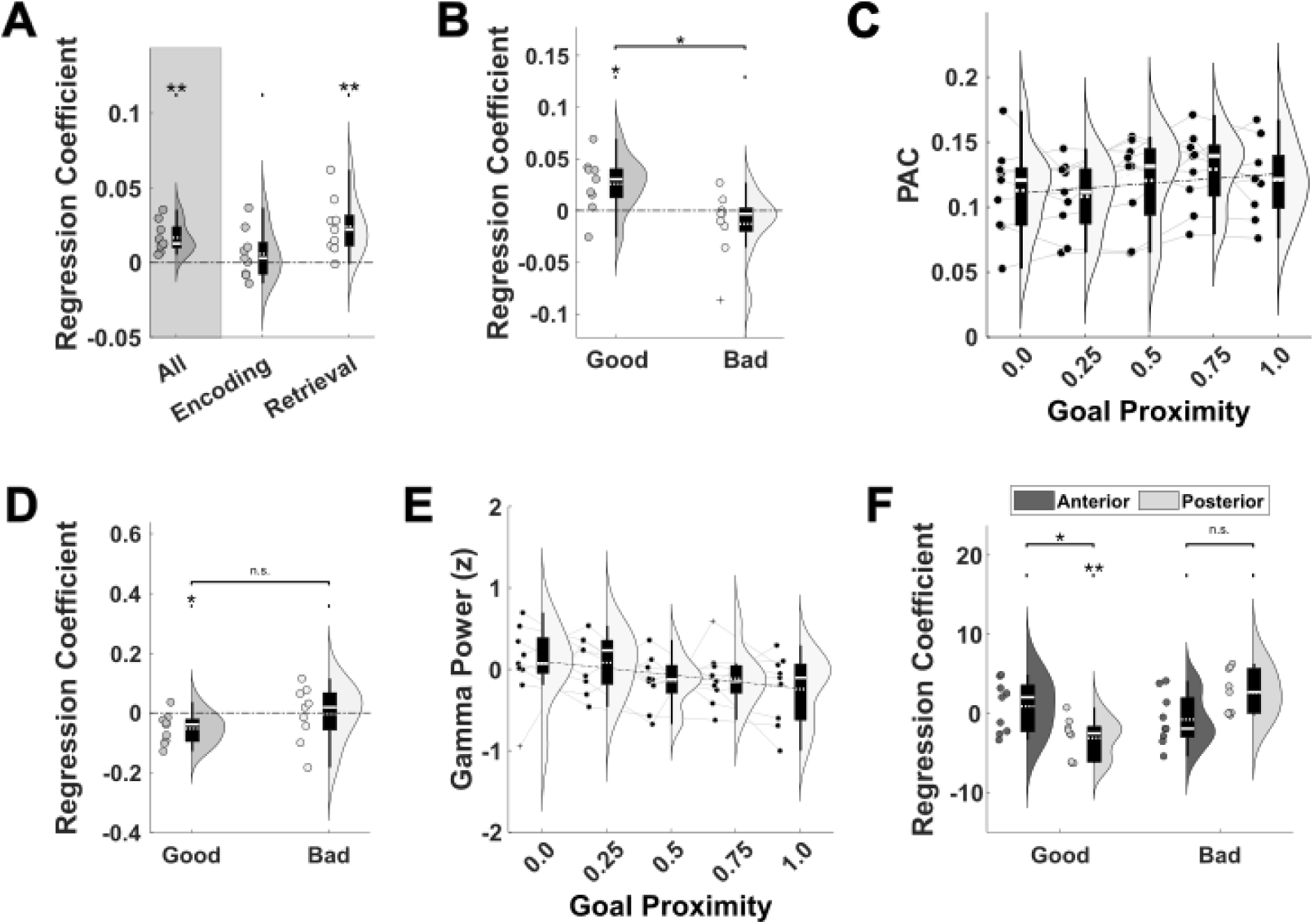
Changes in hippocampal theta-gamma phase-amplitude coupling with goal proximity during memory guided navigation. **(A)** 2-5Hz theta -70-140Hz gamma PAC increases with proximity to the goal location during movement across all trials (t(8) = 4.84, p = 0.00129), but this is driven by memory-guided navigation during retrieval (t(8) = 3.80, p = 0.005, one-sample t-test) rather than visually-guided navigation during encoding (t(8) = 1.05, p = 0.321), although there was no significant difference between encoding and retrieval trials (t(8) = -1.86, p = 0.1). **(B)** The TG-PAC code for goal distance is stronger in good trials (t(8) = 2.84, *p* = 0.0216) than bad trials (t(8) = -1.19, *p* = 0.269; contrast: t(8) = 2.76, *p* = 0.0244). **(C)** Change in TG-PAC with proximity to the goal across all hippocampal electrode contacts during ’good’ retrieval trials. **(D)** 70-140Hz gamma power decreases with proximity to the goal during ’good’ (t(8) = -3.05, *p* = 0.0158), but not ’bad’ retrieval trials (t(8) = -0.115, *p* = 0.912), although there was no significant difference between good and bad trials (t(8) = -1.55, *p* = 0.159). **(E)** Change in gamma power with proximity to the goal across all hippocampal electrode contacts during ’good’ retrieval trials. **(F)** Change in the preferred theta phase of 70-140Hz gamma power with goal proximity during memory guided navigation. There is a significant, negative circular-linear correlation during good trials on posterior electrode contacts (t(8) = 3.69, *p* < 0.001), and a significant difference between regression coefficients in good and bad trials on posterior electrode contacts (t(8) = -4.60, p = 0.0018)

Next, we asked whether the relationship between TG-PAC and distance to the remembered goal location during retrieval trials differed according to accuracy or between anterior and posterior hippocampal recording sites, where the scale of spatial representations may differ. To do so, we entered regression coefficients into a 2 x 2 repeated measures ANOVA with factors of memory performance (good vs bad) and recording location (anterior vs posterior). This showed a main effect of memory performance (F(1,8) = 5.96, *p* = 0.04) but no other main effect or interaction (both p > 0.76). Post-hoc tests indicated that this was driven by the TG-PAC code for goal distance being stronger in good trials (t(8) = 2.84, *p* = 0.0216, one-sample t-test) than bad trials (t(8) = -1.19, *p* = 0.269; contrast: t(8) = 2.76, *p* = 0.0244, paired t-test; Figures 3B, C).

Similar effects were observed for cross-frequency coupling between 2-5 Hz theta and 30 – 70 Hz low gamma (Supplementary Figure 1). Importantly, these effects were not driven by changes in theta waveform asymmetry, which is not modulated by distance from the goal (Supplementary Figure 2); by concomitant changes in theta or gamma power (explored in more detail below), which might bias estimates of phase-amplitude coupling (Supplementary Figure 3); or by distance from the boundaries of the environment, which was strongly (negatively) correlated with distance from the goal location across trials (Supplementary Figure 4). Finally, we asked whether the relationship between TG-PAC and distance to the goal during memory guided navigation is restricted to the hippocampus, or present across temporal lobe recording sites (see Supplementary Tables 3 and 4 for further details). However, we found no evidence for a similar relationship on electrode contacts implanted in either the amygdala or temporal neocortex, in either the low- or high-gamma band (all p > 0.08; Supplementary Figure 5).

In sum, these results indicate that TG-PAC across the long-axis of the hippocampus increases dynamically as the goal is approached during translational movement in a virtual environment, consistent with previous MEG data from a more abstract navigation task (Huang et al., 2026). Moreover, this effect is specific to retrieval trials, when the goal location is hidden and navigation is memory-guided; and is stronger during accurate spatial navigation.

### Hippocampal Gamma Power Decreases with Proximity to the Goal

Next, we investigated whether increases in theta-gamma phase-amplitude coupling with proximity to the remembered goal location are accompanied by reductions in gamma power, as predicted by our theoretical framework (see Figure 2). To examine this, we focussed on 70-140Hz high gamma power during ‘good’ retrieval trials, where the relationship between TG-PAC and goal distance was most pronounced. Consistent with our prediction, we observed a significant decrease in 70-140Hz gamma power with proximity to the hidden goal location (t(8) = -3.05, *p* = 0.0158, one-sample t-test), which was not observed during ‘bad’ trials (t(8) = -0.115, *p* = 0.912, one-sample t-test), although the difference between good and bad trials was not significant (t(8) = -1.55, *p* = 0.159, paired t-test; Figure 3D).

Subsequent analysis indicated that this effect was present on both anterior (t(8) = -2.67, *p* = 0.0283, one-sample t-test) and posterior (t(8) = -3.12, *p* = 0.0142, one-sample t-test) hippocampal electrodes and is significantly stronger in good trials than bad trials in the posterior hippocampus (t(8) = -2.40, *p* = 0.0431, one-sample t-test). However, this effect was not present during encoding trials (t(9) = -0.653, *p* = 0.532, one-sample t-test), or during ‘good’ retrieval trials on electrode contacts in either the amygdala or temporal neocortex (both p > 0.24). We observed the same pattern of results for the relationship between low gamma power and distance to the hidden goal location, although this did not reach significance (Supplementary Figure 6). In sum, these findings suggest that hippocampal high gamma power decreases as a hidden goal location is approached during memory-guided navigation, consistent with our hypothesis that this process is supported by hippocampal theta-gamma phase-amplitude coupling.

### Theta Phase Precession of Gamma Power during Navigation to a Hidden Goal

Next, we asked if the preferred theta phase of high gamma power changed with proximity to the remembered goal location during retrieval trials (see Figure 2). To do so, we quantified the circular-linear relationship between theta phase weighted by gamma amplitude and normalised distance to the goal. Again, we began by focussing on 70-140Hz high gamma power during ‘good’ retrieval trials, where the relationship between TG-PAC and goal distance was most pronounced. Across all electrode contacts, however, we found no consistent relationship between preferred phase and proximity to the remembered goal location (t(8) = -0.84, *p* = 0.421, one-sample t-test).

Nonetheless, we entered circular-linear regression coefficients into a 2 x 2 repeated measures ANOVA with factors of memory performance (good vs bad) and recording location (anterior vs posterior), to ascertain whether the effect might be specific to posterior hippocampal electrode channels, where hippocampal place fields are typically smaller and the spatial representation therefore changes more rapidly (Kjelstrup et al., 2008; Liu et al., 2023). This revealed a significant interaction between recording location and performance (F(1,8) = 124.2, *p* < 0.001) but no main effect of location or performance alone (both *p* > 0.15). Post-hoc tests indicated that this was driven by a significant difference between good and bad trials on posterior (t(8) = -4.60, *p* = 0.0018, paired t-test) but not anterior (t(8) = 1.03, *p* = 0.332, paired t-test) electrode contacts. Crucially, this arose from a significant negative circular-linear correlation between the preferred theta phase of gamma amplitude and normalised distance from the remembered goal location in ‘good’ trials on posterior electrode contacts (t(8) = -3.69, *p* = 0.0062, one-sample t-test; Figure 3F).

In sum, these findings suggest that high gamma power in the posterior hippocampus peaks at progressively earlier theta phases as a hidden goal location is approached during memory-guided navigation, analogous to the phase precession of place cell firing observed in rodent hippocampus (O’Keefe and Recce, 1993) and consistent with our hypothesis that this process is reflected in hippocampal theta-gamma phase-amplitude coupling.

### No Change in Theta Power with Distance to the Goal

Finally, we asked whether these effects might be accompanied by changes in hippocampal theta power. We have previously shown that theta power at movement onset is greater prior to longer spatial trajectories in these data, without considering the impact of distance to the goal location during navigation (Bush et al., 2017). Elsewhere, movement related theta power has been shown to correlate with goal distance when navigating between discrete locations using both MEG (Huang et al., 2026) and intracranial EEG (Liu et al., 2023). However, we did not find any evidence for a relationship between goal proximity and 2-5Hz hippocampal theta power here, either across all trials (t(8) = -0.721, *p* = 0.491, one-sample t-test), or during encoding (t(8) = -0.50, *p* = 0.63, one-sample t-test) and retrieval (t(8) = 0.029, *p* = 0.977, one-sample t-test) trials when analysed separately. Focusing on retrieval trials, a 2 x 2 repeated measures ANOVA with factors of memory performance (good vs bad) and recording location (anterior vs posterior) also showed no main effect or interaction (all *p* > 0.38). Similar results were observed in the 6-9Hz high theta band (all *p* > 0.28), in the amygdala (all *p* > 0.36) and temporal neocortex (all *p* > 0.1). Surprisingly, this suggests that there is no relationship between hippocampal theta power during navigation and the distance to a visible or hidden goal location, in contrast to previous reports tasks (Liu et al., 2023; Huang et al., 2026).

## Discussion

We have shown that theta-gamma phase-amplitude coupling in the human hippocampus encodes the distance to hidden goal locations during memory-guided navigation. This is consistent with the hypothesis that gamma activity reflects the activity of underlying place cell assemblies organised into goal-directed theta sweeps, as described by recent rodent studies (Wikenheiser et al., 2015; Tang et al., 2026; Yu et al., 2026). As the hidden goal location is approached, and the sequence of locations remaining to be traversed becomes shorter, this predicts an increase in theta-gamma phase-amplitude coupling, a decrease in gamma power, and the theta phase precession of gamma power. We have verified each of these predictions in direct intracranial recordings of the hippocampal local field potential (LFP) in people with epilepsy performing a naturalistic virtual navigation task. These results extend our previous findings from MEG recordings of healthy participants performing an abstract item navigation task (Huang et al., 2026), replicating those results using a more ethologically valid spatial memory paradigm and without relying on the source localisation of high-frequency activity from deep structures, which may be problematic (but see Gohil et al., 2026). More generally, these findings are compatible with the hypothesis that the hippocampus supports model - based roll-out across species and task domains, evaluating different sequences of possible actions to support efficient memory-guided behaviour (Daw et al., 2005; Behrens et al., 2018).

These findings build on an extensive theoretical and empirical literature implicating theta-gamma phase-amplitude coupling in working memory maintenance (Lisman and Idiart, 1995; Jensen and Lisman, 1998; Mormann et al., 2005; Axmacher et al., 2010; Lisman and Jensen, 2013) and, more recently, in sequence learning (Heusser et al., 2016). Consistent with our results, cross-frequency coupling has been shown to decrease (Daume et al., 2024) and gamma power has been shown to increase (Howard et al., 2003; Van Vugt et al., 2010) with working memory load. Similarly, theta-gamma phase-amplitude coupling has been shown to decrease with the length of a sequence being encoded into long-term memory (Heusser et al., 2016), as gamma activity become more widely spread across the theta cycle. In each case, this coding scheme supports the multiplexing of stimulus features and ordinal structure for a limited number of items. The key distinction between our results and previous findings is that this ordinal structure is endogenously generated during sequential planning in the service of goal-directed navigation, in contrast to the passive encoding of externally presented sequences in working and episodic memory paradigms.

Importantly, we did not find any evidence for a relationship between theta power and the distance to hidden or visible goal locations in these data, despite previous evidence for such a signature both in MEG (Huang et al., 2026) and in an intracranial EEG study using a very similar task (Liu et al., 2023). Similarly, we found no evidence for theta-gamma coding of the distance to visible goal locations during encoding trials. In contrast, recent rodent data shows theta sweeps towards dangled and scattered food rewards in the absence of a memory component (Vollan et al., 2026). It is also unclear from the existing rodent data whether theta sweeps terminate at hidden goal locations during memory guided navigation, as we assume here, although it has been demonstrated that the spatial extent of theta sweep trajectories becomes shorter as goal locations are approached (Wikenheiser et al., 2015; Yu et al., 2026). Re-analysis of existing rodent electrophysiology data sets could address this issue, and whether the spatial extent of theta sweeps within each cycle shows the predicted relationship with theta-gamma phase-amplitude coupling (e.g. longer theta sweeps being associated with lower coupling).

In sum, we have shown that theta-gamma phase-amplitude coupling in the human hippocampus encodes the distance to hidden goal locations during spatial navigation. Alongside previous theoretical and experimental findings, this suggests that theta-gamma phase-amplitude coupling may reflect a domain general mechanism of model-based planning.

## Materials and Methods

### Patient Population

The data analysed here were previously described in Bush et al. (2017). Briefly, nineteen patients with pharmaco-resistant epilepsy who underwent intracranial EEG to determine surgical candidacy completed a spatial memory task in desktop virtual reality (VR). Three patients were excluded due to technical difficulties, six had no electrode contacts inside the hippocampus, and one could not perform the task. This left nine patients: five female and seven right-handed, with a mean ± SD age of 30.8 ± 7.9 years. All patients had platinum Spencer probe depth electrodes (Ad-Tech Medical) targeted at the medial temporal lobe: seven in the right hemisphere, one in the left hemisphere, and one bilateral implantation. This study was approved by the NHS research ethics committee, and all patients gave written informed consent. Patients were seizure-free for at least 24 h before participation.

### Spatial Memory Task

Participants completed a spatial memory task that had been used in several previous neuroimaging studies (Doeller et al., 2008; Kaplan et al., 2012; Kaplan et al., 2014; Adams et al., 2020). In this task, participants are asked to encode and then retrieve the locations of different objects in a 100 virtual metre (vm) sided square VR environment displayed on a laptop screen in front of them. Participants can navigate freely throughout the task using the arrow keys to move forwards and rotate left or right, with movement accelerating rapidly to a fixed top speed of 30 vm/s. Their location and orientation within the VR environment were recorded continuously throughout all trials at a sample rate of 25 Hz.

During encoding, participants are asked to encode the locations of four different visible objects presented in pseudo-random order across five blocks, giving a total of 20 encoding trials. In each encoding trial, they navigate freely around the environment, and when they are satisfied that they have encoded the location of the visible object, they collide with that object to proceed to the next trial. Participants are then given a 30 s break, during which they are instructed to focus on memorising object locations.

During retrieval, participants are cued with an image of one object, placed back into the environment at one of four randomly selected locations, and instructed to navigate to the remembered location of that object and make a response using the space bar. After responding, the object appears in its correct location to provide feedback, and participants are instructed to collide with the visible object to proceed to the next trial. Each retrieval begins with a 3 s fixation period, followed by the target object image presented for 3 s on a plain grey background; and there are five retrieval trials for each object presented in a pseudorandom order, giving a total of 20 retrieval trials.

Three participants completed one session of the task and six participants completed two sessions. The second session used the same VR environment but used four new objects at different locations. To quantify task performance, the error *Δd* in each trial was defined as the Euclidean distance between the response location and true object location. Distance errors were then normalised relative to the distribution of all possible distance errors, such that 1 is the best possible performance, 0 is the worst possible performance, and 0.5 is chance performance for each object (Miller et al., 2018). Good and bad trials were defined as those above and below the median of normalised distance errors within each session, respectively.

Finally, we define the ‘remembered goal location’ as the response location in each retrieval trial. Distance from the visible object location during encoding, true and remembered goal locations during retrieval were also normalised relative to the distribution of all possible Euclidean distances from those locations across the whole environment, subtracted from 1. As such, 0 indicates the furthest possible distance from the visible, true, or remembered object location, and 1 indicates arrival at that location. Similarly, we define the ‘boundary distance’ as the Euclidean distance between the participant’s current location and the closest boundary. Boundary distances were also normalised, such that 1 indicates a location directly adjacent to the boundary, and 0 indicates a location in the centre of the environment.

### Intracranial EEG Recordings

Intracranial EEG was recorded continuously throughout the task at a sample rate of 1024 Hz (patients 4 and 8) or 512 Hz (all other patients) using either a Nicolet NicoletOne long-term monitoring system (Natus Medical, Inc.; patients 1-5) or a Micromed SD long-term monitoring system (Micromed; all other patients). To align neural and behavioural data, audio triggers produced by the VR task were recorded alongside the intracranial EEG signal. Prior to any analyses, EEG signals from all channels and recording sessions were visually inspected for interictal spike (IIS) events and other artefacts by a trained epileptologist (UV) and verified by an EEG analysis expert (DB).

Electrode locations were determined solely by clinical criteria and confirmed by visual inspection of post-implantation CT and/or MRI scans. The presence of hippocampal damage was also assessed, when histopathology was not available, by structural MRI at 3T including quantification of hippocampal volumes and T2 relaxation times. Our analyses focussed on EEG recordings from three different regions of interest: the hippocampus (nine patients with mean ± SD of 5 ± 1.2 contacts; range of 4-8 contacts); temporal neocortex (nine patients, 8.8 ± 1.5 contacts; range of 6-12 contacts); and amygdala (eight patients, 2.4 ± 0.7 contacts; range of 2-3 contacts).

### Epochs of Interest

Our analyses focussed on periods of continuous translational movement throughout encoding and retrieval trials, up to the point that a response was made (e.g. excluding feedback periods). We defined movement epochs as any period of continuous translation across the environment with speed >1 m/s that last >1 s. Movement epochs were subsequently segmented into non-overlapping 1 s windows, and the normalised distance at the start of each epoch from the visible, true and / or remembered goal location was computed for each sliding window during encoding and retrieval trials, respectively. Sliding windows with IIS occurring within 0.5s of the start or end were excluded from all subsequent analysis.

### Time Frequency Analysis

Estimates of dynamic oscillatory power during each epoch of interest were generated using MNE (Gramfort et al., 2013). Specifically, power estimates were obtained for 50 logarithmically spaced frequency bands in the 2-150 Hz range by convolving the EEG signal with a five-cycle Morlet wavelet. Power values were log-transformed but not baseline corrected. For movement epochs, time-frequency representations were generated for windows beginning 2 s before and ending 2 s after the onset and offset of each sliding window, respectively, before being cropped to the 1 s sliding window to avoid edge effects. For cue periods, time-frequency representations were generated for windows beginning 2 s before and ending 2 s after cue onset and offset, respectively, before being cropped to the 3 s window of interest.

### Phase-amplitude Coupling

To examine cross-frequency coupling between theta phase and gamma amplitude, we first filtered the signal in each frequency band of interest using a fourth-order zero-phase Butterworth filter. Dynamic estimates of phase and amplitude in each frequency band were then obtained using the Hilbert transform. Finally, phase-amplitude coupling (PAC) was quantified as the mean vector length of theta phases weighted by instantaneous gamma amplitude across the time window of interest (Canolty et al., 2006). We focussed our analyses on the 2-5Hz ‘low’ theta band from our previous MEG (Huang et al., 2026) and intracranial EEG (Bush et al., 2017) studies, and on the 70-140Hz ‘fast’ and 30-70Hz ‘slow’ gamma band from our previous MEG study (Huang et al., 2026).

To quantify whether PAC covaries with distance from the goal during movement, we computed linear regression coefficients between PAC estimates and normalised distance to the visible location (during encoding trials), true or remembered goal location (during retrieval trials) across all 1 s sliding windows on each electrode contact, with trial number included as a nuisance regressor. Average regression coefficients across all sessions and channels were then submitted to a one-sample t-test across participants.

To examine whether changes in phase-amplitude coupling are driven by concurrent changes in power in the theta and gamma frequency bands, we performed a linear regression between the trial-by-trial PAC estimates and the theta and gamma power. We took the residuals after regressing out the power effect and conducted the same linear regression analysis between the PAC residuals and the goal proximity.

To further examine whether changes in PAC as the goal is approached are driven by the influence of distance to the nearest boundary, we included both goal proximity and the distance to the boundary as regressors in the regression analysis. For each participant, we obtain one regression coefficient average across channels and sessions for each regressor. Average regression coefficients across all sessions and channels were then submitted to a one-sample t-test across participants.

### Theta Waveform Asymmetry

To quantify theta waveform asymmetry, we filtered the signal in the 2-5 Hz theta band using a fourth order zero-phase Butterworth filter. Local minima and maxima of the filtered signal in each non-overlapping 1 s window were then obtained and used to calculate the asymmetry index, which is defined as the difference between the ascending (peak to trough) and descending (trough to phase) phases of the theta cycle, normalised by the length of the theta cycle (Belluscio et al., 2012). Finally, these values were entered into the same regression analysis with normalised distance to the visible location (during encoding trials), true or remembered goal location (during retrieval trials) across all 1 s sliding windows on each electrode contact, with trial number included as a nuisance regressor. Average regression coefficients across all sessions and channels were then submitted to a one-sample t-test across participants.

### Preferred Theta Phase of Gamma Power

To investigate whether the preferred 2-5 Hz theta phase of 70–140 Hz gamma power covaries with distance from the goal during movement, we first computed the circular mean of theta phases weighted by instantaneous gamma amplitude for each 1 s non-overlapping window during movement. Next, we calculated the circular-linear correlation (Kempter et al., 2012) between preferred theta phases and normalised distance to the visible location (during encoding trials), true or remembered goal location (during retrieval trials) across all sliding windows on each electrode contact. Average circular-linear slopes across all sessions and channels were then submitted to a one-sample t-test across participants.

All analyses were conducted using custom-written Python code, with circular statistics computed using the PyCircStat2 toolbox (https://circstat.github.io/pycircstat2/).

## Acknowledgements

This work was supported by the Department of Health’s National Institute for Health Research, UCL/UCL Biomedical Research Centre. U.V. is funded by a Wellcome Trust Career Development Award (307306/Z/23/Z), the Medical Research Foundation and NIHR BRC. N.B. is funded by a Wellcome Trust Principal Research Fellowship (222457/Z/21/Z). D.B. is funded by a UKRI Frontier Research Grant (EP/X023060/1).

## Supplementary Information

**Supplementary Figure 1:**
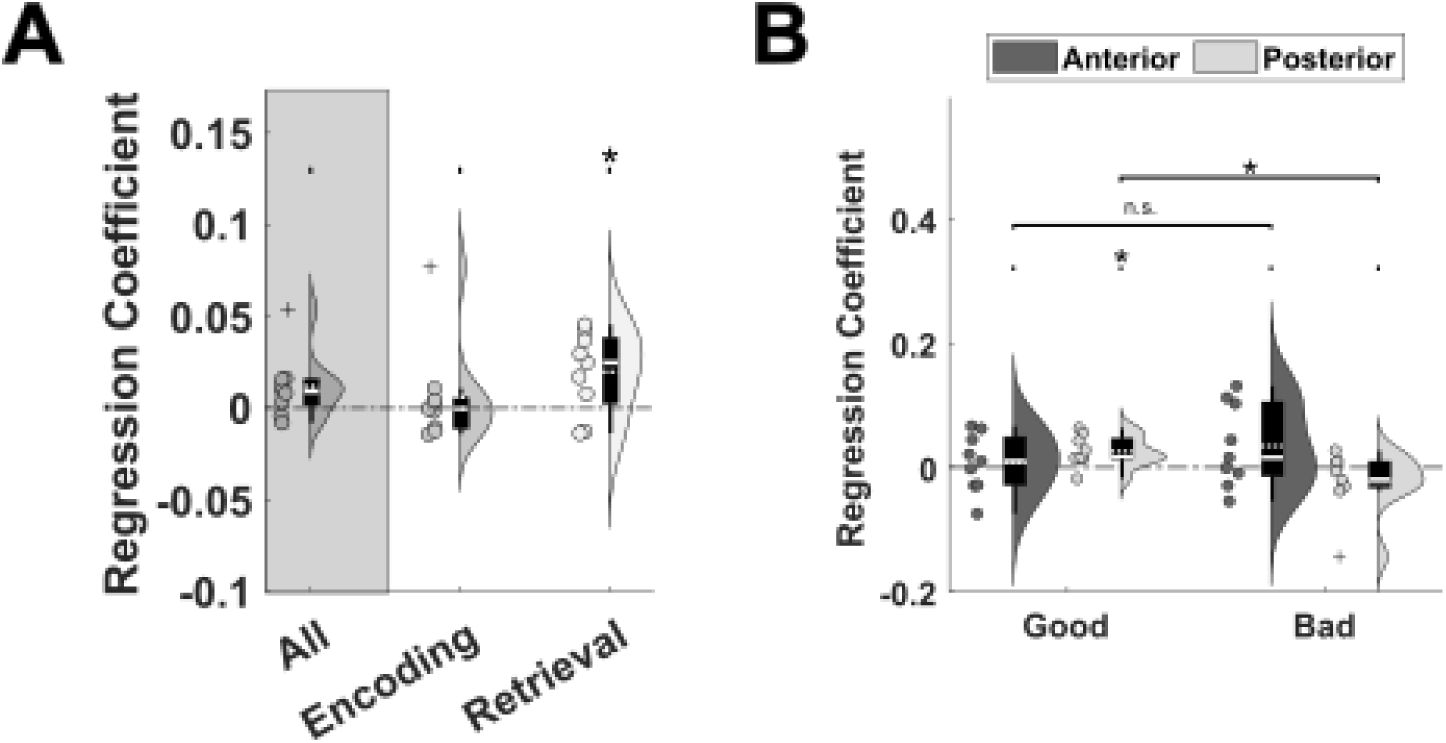
Theta – low gamma PAC is also correlated with distance to hidden goal locations. **(A)** Across both encoding and retrieval, we found a trend towards TG-PAC increasing with proximity to the (visible and remembered) goal location (t(8) = 1.97, *p* = 0.0838). Again, this effect was driven by memory guided navigation during retrieval (t(8) = 2.66, *p* = 0.0287, one-sample t-test), but not visually guided navigation during encoding (t(8) = 0.660, *p* = 0.122, one-sample t-test), although there was no significant difference between encoding and retrieval trials (t(8) = -1.32, *p* = 0.221, paired t-test). **(B)** A 2 x 2 repeated measures ANOVA revealed no main effect of recording site or memory accuracy during retrieval trials (both *p* > 0.23), although we note a trend towards significance for the interaction (F(1,8) = 3.51, *p* = 0.098). Post-hoc tests indicated that this was driven by a stronger relationship in good trials (t(8) = 3.06, *p* = 0.0155, one-sample t-test) than in bad trials (t(8) = -1.48, *p* = 0.179, one-sample t-test) on posterior hippocampal electrode contacts (contrast: t(8) = 2.80, *p* = 0.023, paired t-test). This suggests that the observed increase in TG-PAC as the goal is approached is also present for the 30 – 70Hz low gamma band.

**Supplementary Figure 2:**
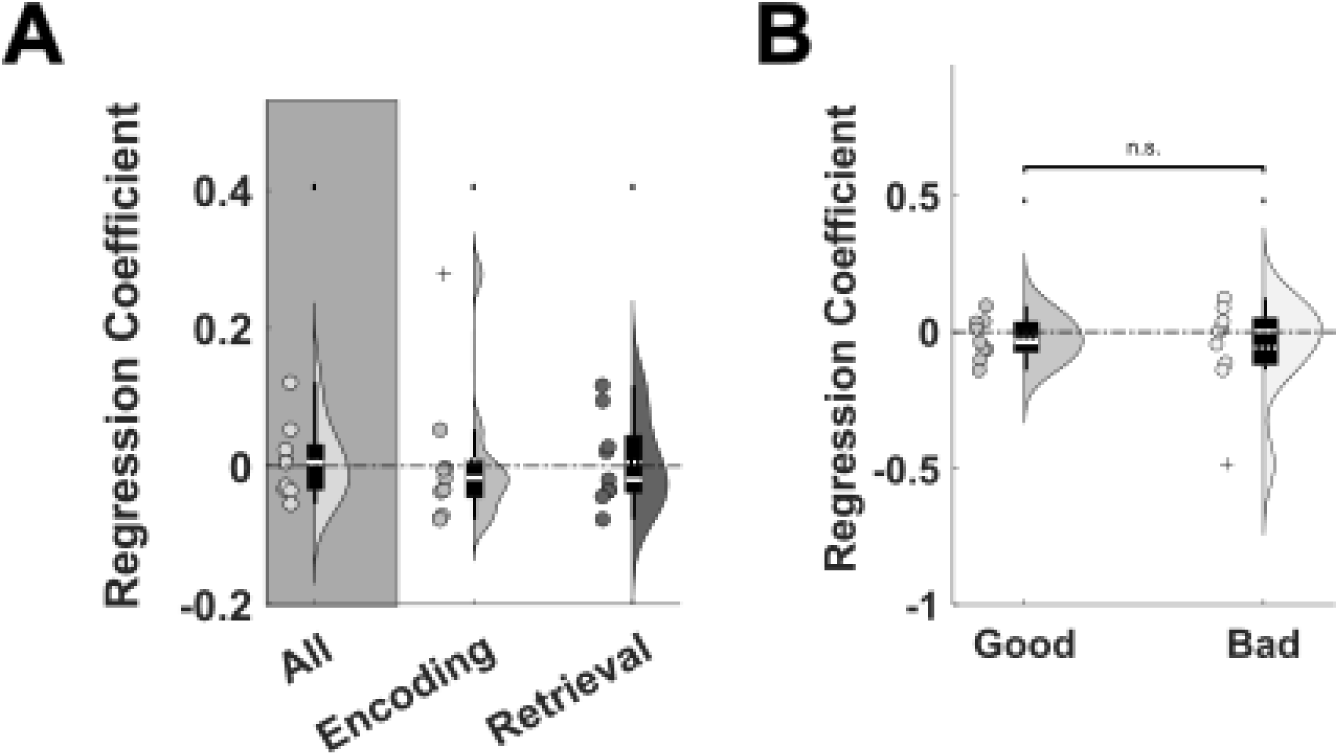
Changes in TG-PAC are not driven by changes in theta waveform asymmetry. To investigate whether the observed changes in TG-PAC with distance to the goal may arise spuriously from changes in theta waveform asymmetry (Scheffer-Teixeira et al., 2016), we calculated the trial-by-trial theta waveform asymmetry index (Belluscio et al., 2012) and entered those values into the same linear regression analyses used for TG-PAC. **(A)** However, we did not find any significant correlation between theta asymmetry and goal proximity during either encoding (t(8) = 0.262, *p* = 0.80, one-sample t-test) or retrieval (t(8) = 0.245, *p* = 0.813, one-sample t-test) trials. **(B)** Similarly, we found no correlation between theta asymmetry and goal proximity if we analysed good (t(8) = -1.01, *p* = 0.341, one-sample t-test) and bad (t(8) = -0.952, *p* = 0.370, one-sample t-test) retrieval trials separately. This suggests that the observed increase in TG-PAC as the goal is approached does not arise spuriously from changes in theta waveform asymmetry.

**Supplementary Figure 3:**
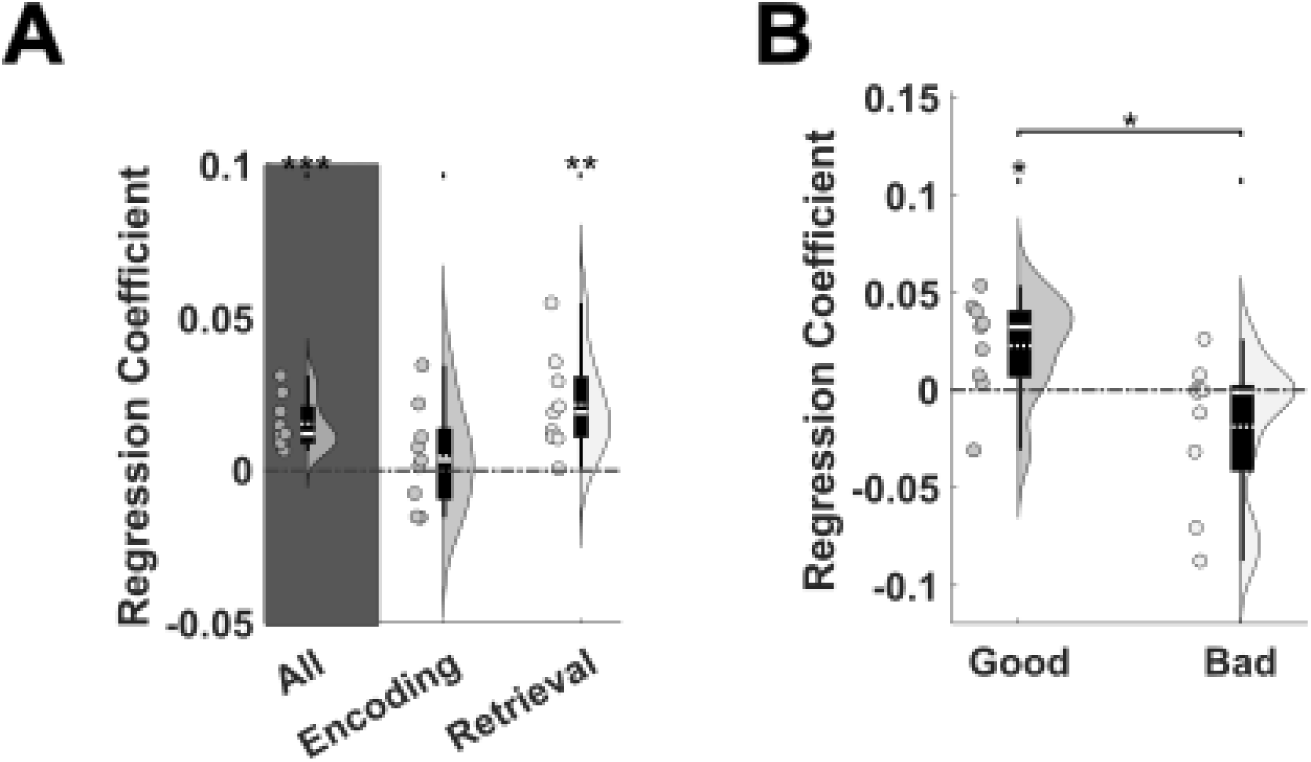
Changes in TG-PAC are not driven by changes in theta or gamma power. To remove any potential confound caused by concurrent changes in oscillatory power, we repeated our linear regression analysis between distance to the goal and TG-PAC estimates while including theta and gamma band power in each sliding window as nuisance regressors. This did not qualitatively affect any of our main results. **(A)** Specifically, we still observed a significant relationship between TG-PAC and distance to the goal across all trials (t(8) = 5.44, *p* < 0.001, one-sample t-test) that was primarily driven by retrieval (t(8) = 3.97, *p* = 0.004, one-sample t-test) but not encoding (t(8) = 0.829, *p* = 0.431, one-sample t-test) trials, although there was no significant difference between task periods (t(8) = -1.76, *p* = 0.116, one-sample t-test). **(B)** Again, this relationship was significantly stronger in good (t(8) = 2.65, *p* = 0.0291) versus bad (t(8) = -1.47, *p* = 0.181, one-sample t-test) retrieval trials (contrast: t(8) = 3.02, *p* = 0.0165, paired t-test). This suggests that the observed increase in TG-PAC as the goal is approached does not arise spuriously from concurrent changes in theta or gamma power.

**Supplementary Figure 4:**
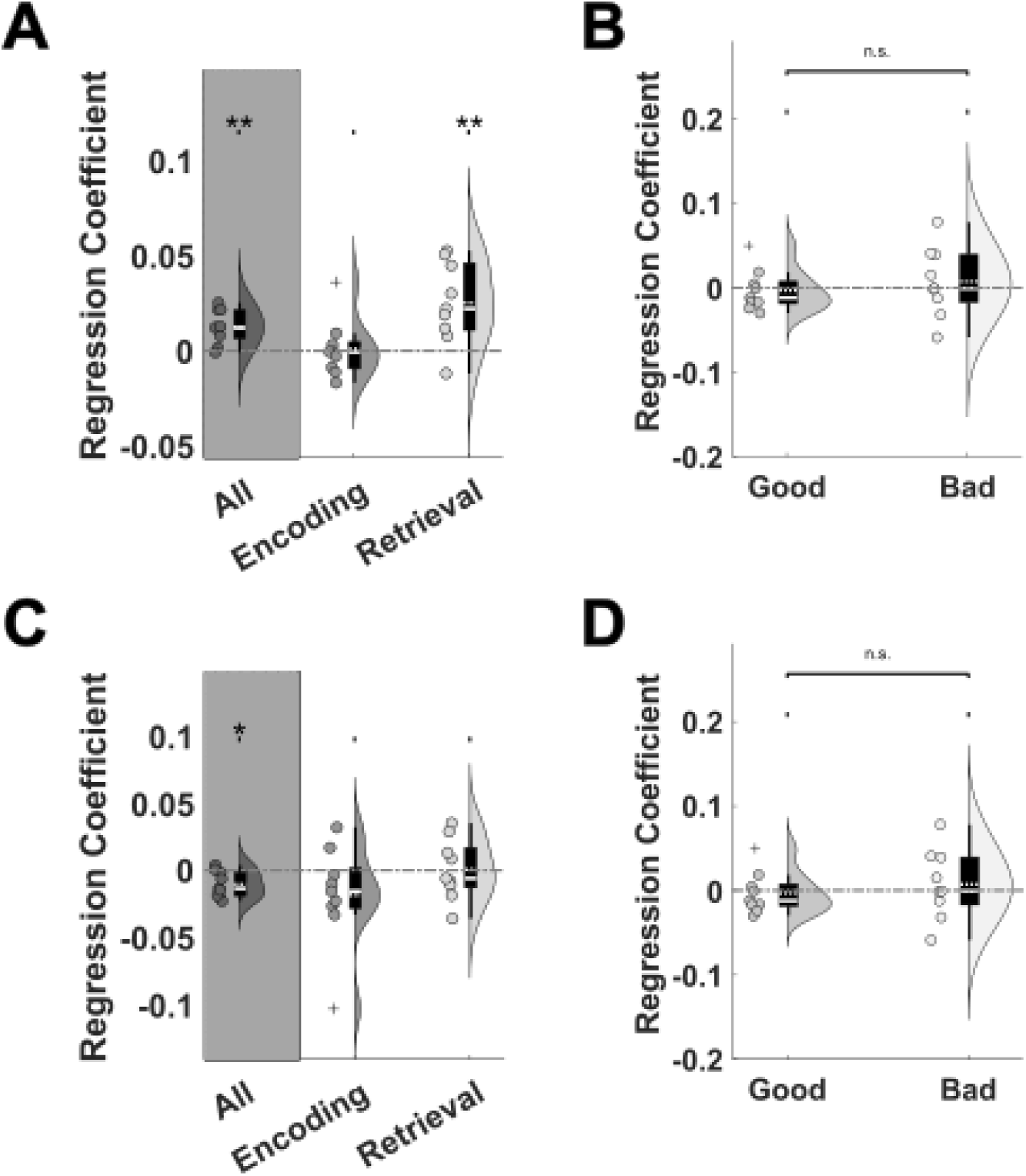
Changes in TG-PAC are not driven by the distance to environmental boundaries. Because distance to the goal was strongly (negatively) correlated with distance to the boundary across all trials (t(8) = -23.5, *p* < 0.001, one-sample t-test), and because boundary proximity has previously been shown to modulate human hippocampal theta power (Stangl et al., 2020), we repeated our linear regression analysis between distance to the goal and TG-PAC estimates while including normalised distance to the closest environmental boundary in each sliding window as a nuisance regressor. This did not qualitatively affect any of our main results. **(A)** Specifically, we still observed a significant correlation between TG-PAC and goal proximity across all trials (t(8) = 4.22, *p* = 0.00293, one-sample t-test) that was significantly stronger during retrieval (t(8) = 3.51, *p* = 0.00795, one-sample t-test) than during encoding (t(8) = 0.170, *p* = 0.869, one-sample t-test; contrast: t(8) = 2.48, *p* = 0.038, paired t-test). **(B)** Again, this relationship was significantly stronger in good (t(8) = 4.17, *p* = 0.00309) versus bad (t(8) = 0.836, *p* = 0.47, one-sample t-test) retrieval trials, although there was no significant difference between good and bad trials (t(8) = 1.06, *p* = 0.317, paired t-test). **(C)** Importantly, although there was a significant negative correlation between distance to the closest boundary and TG-PAC across all trials (t(8) = -3.15, p = 0.0137, one-sample t-test), this did not reach significance during either encoding (t(8) = -1.44, *p* = 0.189, one-sample t-test) or retrieval (t(8) = 0.0109, *p* = 0.92, one-sample t-test) trials in isolation. **(D)** Similarly, we found no correlation between theta asymmetry and goal proximity if we analysed good (t(8) = -0.357, *p* = 0.730, one-sample t-test) and bad (t(8) = 0.552, *p* = 0.56, one-sample t-test) retrieval trials separately. This suggests that the observed increase in TG-PAC as the goal is approached does not arise spuriously from boundary proximity encoding.

**Supplementary Figure 5:**
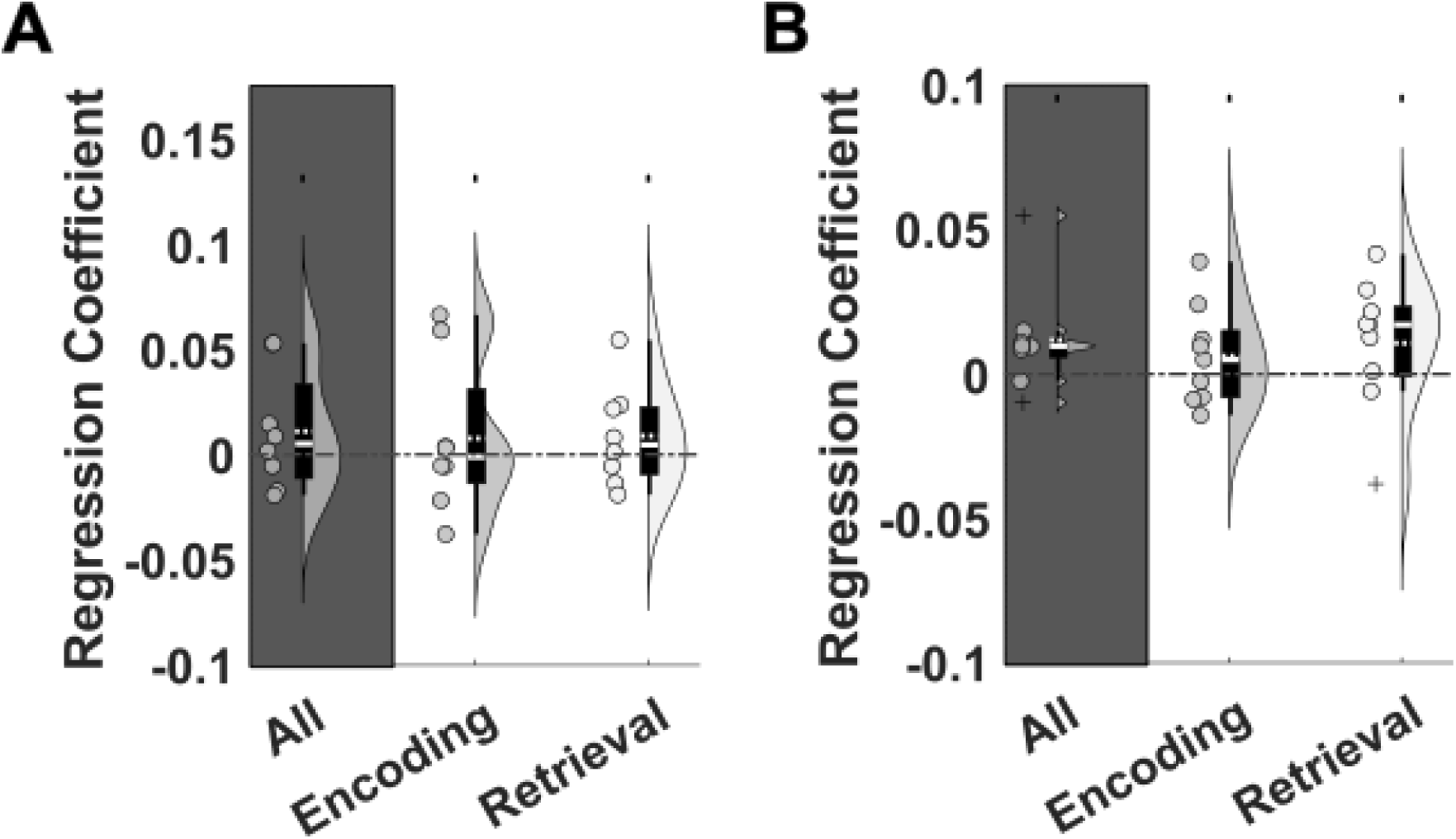
TG-PAC vs distance to the goal in amygdala and temporal neocortex. **(A)** In the amygdala, we found no evidence for a relationship between TG-PAC and goal distance across all trials (t(7) = 1.11, *p* = 0.305), or if we analysed encoding (t(7) = -0.595, *p* = 0.571) and retrieval (t(7) = 1.05, *p* = 0.329) trials separately. Similar results were observed for the relationship between theta – low gamma coupling and goal distance (all *p* > 0.85). **(B)** In addition, we found no evidence for a relationship between TG-PAC and goal distance on electrode contacts in the temporal neocortex, across all trials (t(8) = 1.96, *p* = 0.0844), of if we analysed encoding (t(8) = 1.09, *p* = 0.306) and retrieval (t(8) = 1.39, *p* = 0.201) trials separately. Similar results were observed for the relationship between theta – low gamma coupling and goal distance (all *p* > 0.24). This suggests that the observed increase in TG-PAC as the goal is approached is specific to electrode contacts located in the body of the hippocampus.

**Supplementary Figure 6:**
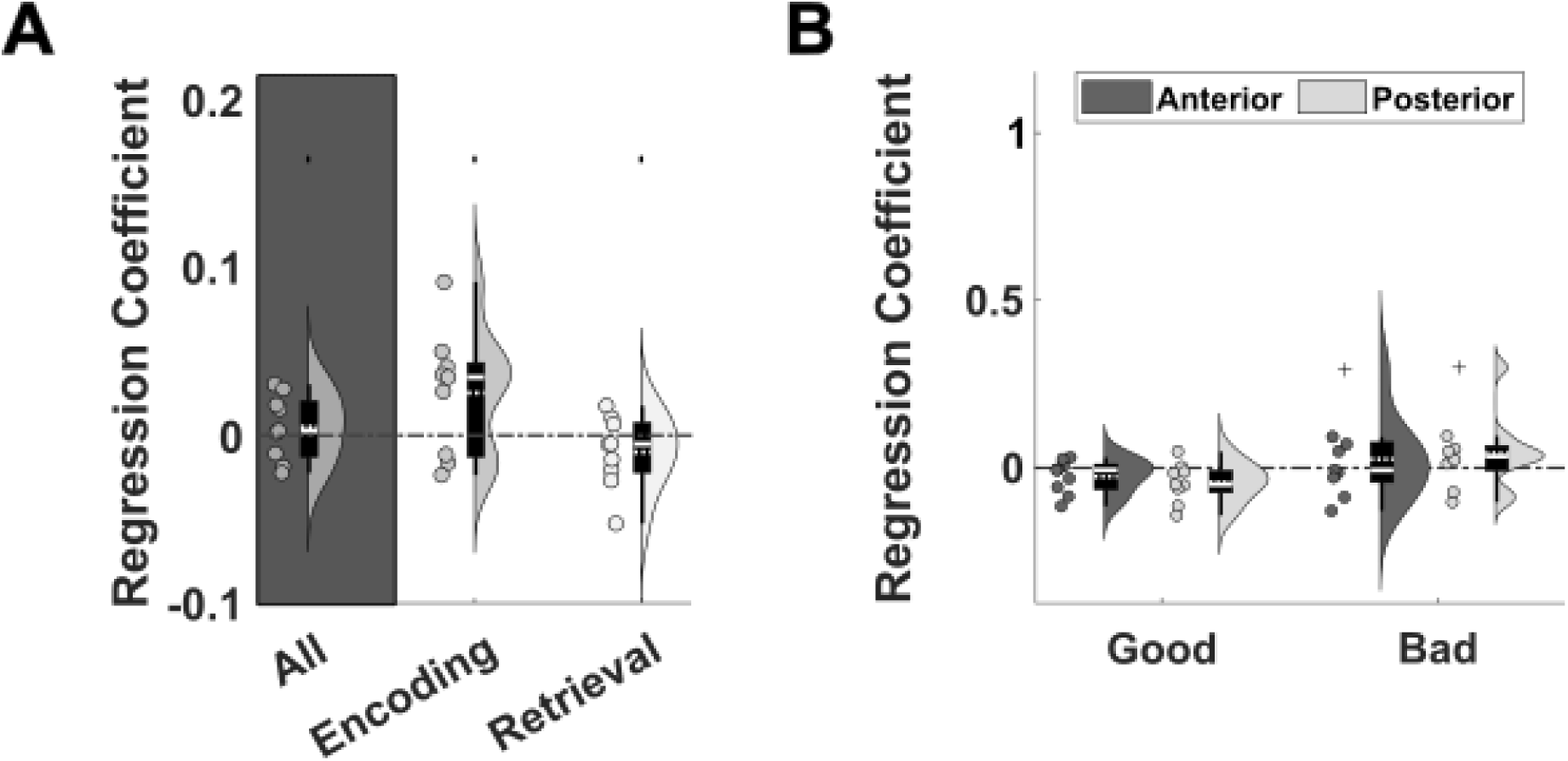
Low gamma power vs distance to the goal in the hippocampus. **(A)** We found no evidence for a relationship between low gamma power and distance to the goal either across all trials (t(8) = 0.746, *p* = 0.477, one-sample t-test), or during encoding (t(8) = 2.05, *p* = 0.0749, one-sample t-test) or retrieval (t(8) = -1.40, *p* = 0.198, one-sample t-test) trials separately. **(B)** Nonetheless, a repeated measures ANOVA focussed on retrieval trials with factors of performance (good vs bad trials) and electrode location (anterior vs posterior hippocampus) showed no main effect of performance or channel position (both *p* > 0.21) but a trend towards an interaction (F(1,8) = 4.70, *p* = 0.062). Post-hoc tests indicated that this was driven by the difference between good and bad trials being greater for posterior vs anterior contacts (t(8) = 2.17, *p* = 0.0621, paired t-test). Specifically, there was a trend towards a negative relationship in good retrieval trials on posterior electrode contacts (t(8) = -2.23, *p* = 0.0563, one-sample t-test), but not in good trials on anterior electrode contacts or bad trials on either anterior or posterior electrode contacts (all *p* > 0.16). This suggests that low gamma power on posterior electrode contacts does increase as the goal is approached during good retrieval trials, although this relationship does not reach significance.

**Supplementary Table 1:** Patient demographics.

| Patient ID | Age (years) | Handed | Sex | Implanted Hemisphere | Sessions Completed | HPC Contacts (ant. / post.) | Amy Contacts | Temp Nctx Contacts |
| --- | --- | --- | --- | --- | --- | --- | --- | --- |
| 1 | 21 | R | F | R | 2 | 4 (3/1) | 0 | 6 |
| 2 | 44 | R | F | R | 2 | 5 (2/3) | 2 | 8 |
| 3 | 31 | R | M | L | 1 | 4 (2/2) | 3 | 9 |
| 4 | 41 | L | M | R | 1 | 5 (3/2) | 2 | 9 |
| 5 | 26 | R | M | R | 2 | 5 (3/2) | 3 | 8 |
| 6 | 37 | R | F | R | 2 | 4 (3/1) | 3 | 10 |
| 7 | 30 | R | F | Bilateral | 2 | 8 (4/4) | 4 | 12 |
| 8 | 20 | L | F | R | 2 | 5 (3/2) | 3 | 9 |
| 9 | 28 | R | M | R | 1 | 5 (3/2) | 2 | 8 |

**Supplementary Table 1:** Patient demographics
| Patient ID | Age at First Seizure | Language Dominance | Seizure Onset Zone | Surgical Outcome | Hippocampal Damage? | Additional Notes |
| --- | --- | --- | --- | --- | --- | --- |
| 1 | 14 yrs | L | Right MTL | Right MTL resection | Right hippocampal sclerosis, severe neuronal loss in CA1 | Right parietal gyral calcification |
| 2 | 23 yrs | L | Right MTL | Right MTL resection | End folium sclerosis of the right hippocampus |  |
| 3 | 18 mo | L | Left temporal | Left TL resection | None | Dysembryoplastic neuroepithelial tumor |
| 4 | 25 yrs | L | Right MTL | Right MTL resection | Right hippocampal sclerosis, severe neuronal loss in CA1 | Periventricular heterotopia |
| 5 | 13 yrs | B | Right neocortical TL | Right neocortical TL resection | None |  |
| 6 | 21 yrs | L | Right temporal-occipital | DNT removal from right temporal-occipital cortex | None |  |
| 7 | 23 yrs | L | Bilateral MTL | No surgery | None | Small left occipito-temporal sulcus lesion |
| 8 | 6 yrs | L | Right TPJ and MTL | Right MTL resection | None |  |
| 9 | 8 yrs | L | Right neocortical TL | No surgery | None | Right temporal / parietal cortical dysplasia, polymicrogyria |

**Supplementary Table 2:**
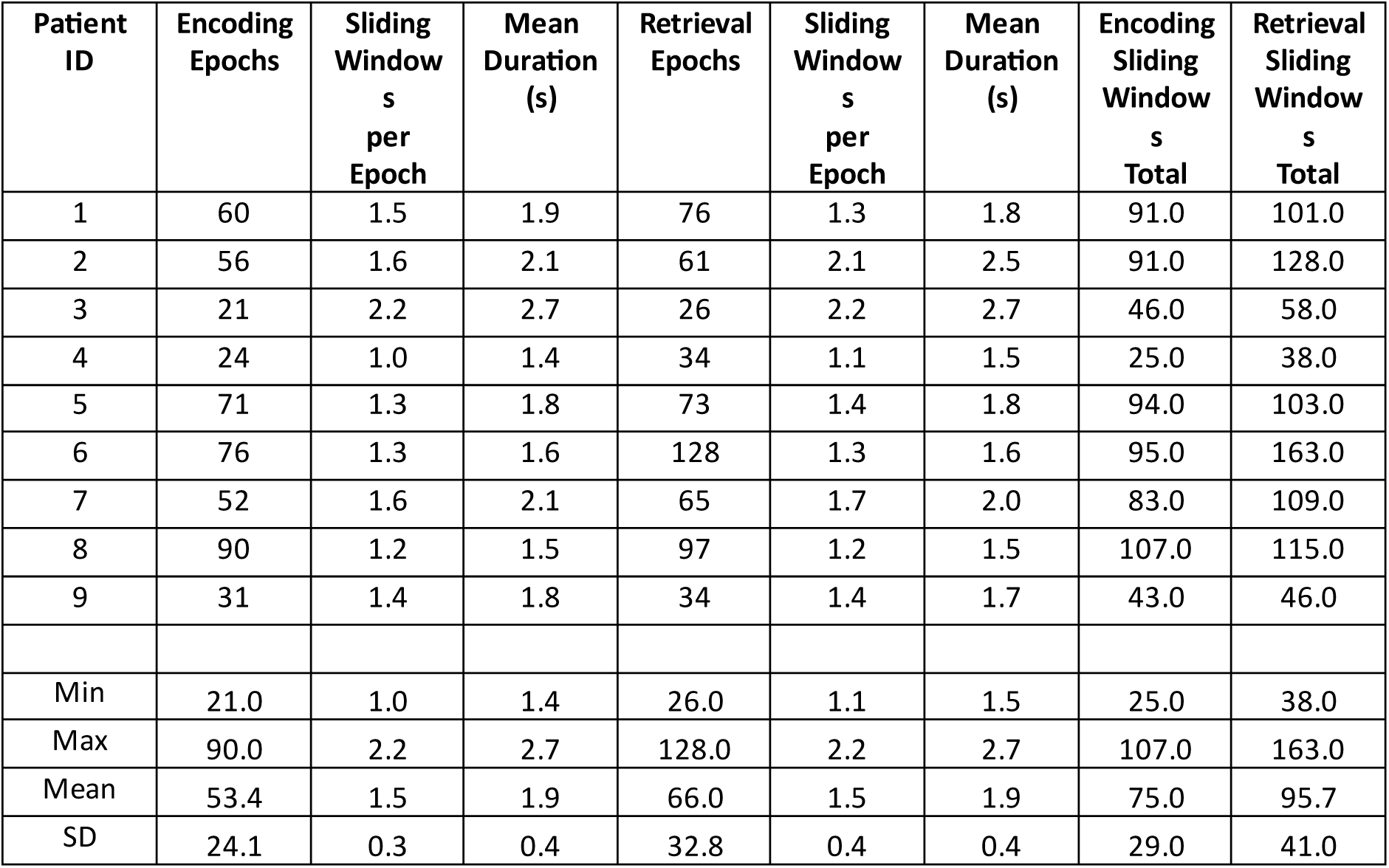
Task epochs for depth electrode contacts in the hippocampus.

| Patient ID | Encoding Epochs | Sliding Windows per Epoch | Mean Duration (s) | Retrieval Epochs | Sliding Windows per Epoch | Mean Duration (s) | Encoding Sliding Windows Total | Retrieval Sliding Windows Total |
| --- | --- | --- | --- | --- | --- | --- | --- | --- |
| 1 | 60 | 1.5 | 1.9 | 76 | 1.3 | 1.8 | 91.0 | 101.0 |
| 2 | 56 | 1.6 | 2.1 | 61 | 2.1 | 2.5 | 91.0 | 128.0 |
| 3 | 21 | 2.2 | 2.7 | 26 | 2.2 | 2.7 | 46.0 | 58.0 |
| 4 | 24 | 1.0 | 1.4 | 34 | 1.1 | 1.5 | 25.0 | 38.0 |
| 5 | 71 | 1.3 | 1.8 | 73 | 1.4 | 1.8 | 94.0 | 103.0 |
| 6 | 76 | 1.3 | 1.6 | 128 | 1.3 | 1.6 | 95.0 | 163.0 |
| 7 | 52 | 1.6 | 2.1 | 65 | 1.7 | 2.0 | 83.0 | 109.0 |
| 8 | 90 | 1.2 | 1.5 | 97 | 1.2 | 1.5 | 107.0 | 115.0 |
| 9 | 31 | 1.4 | 1.8 | 34 | 1.4 | 1.7 | 43.0 | 46.0 |
| Min | 21.0 | 1.0 | 1.4 | 26.0 | 1.1 | 1.5 | 25.0 | 38.0 |
| Max | 90.0 | 2.2 | 2.7 | 128.0 | 2.2 | 2.7 | 107.0 | 163.0 |
| Mean | 53.4 | 1.5 | 1.9 | 66.0 | 1.5 | 1.9 | 75.0 | 95.7 |
| SD | 24.1 | 0.3 | 0.4 | 32.8 | 0.4 | 0.4 | 29.0 | 41.0 |

**Supplementary Table 3:**
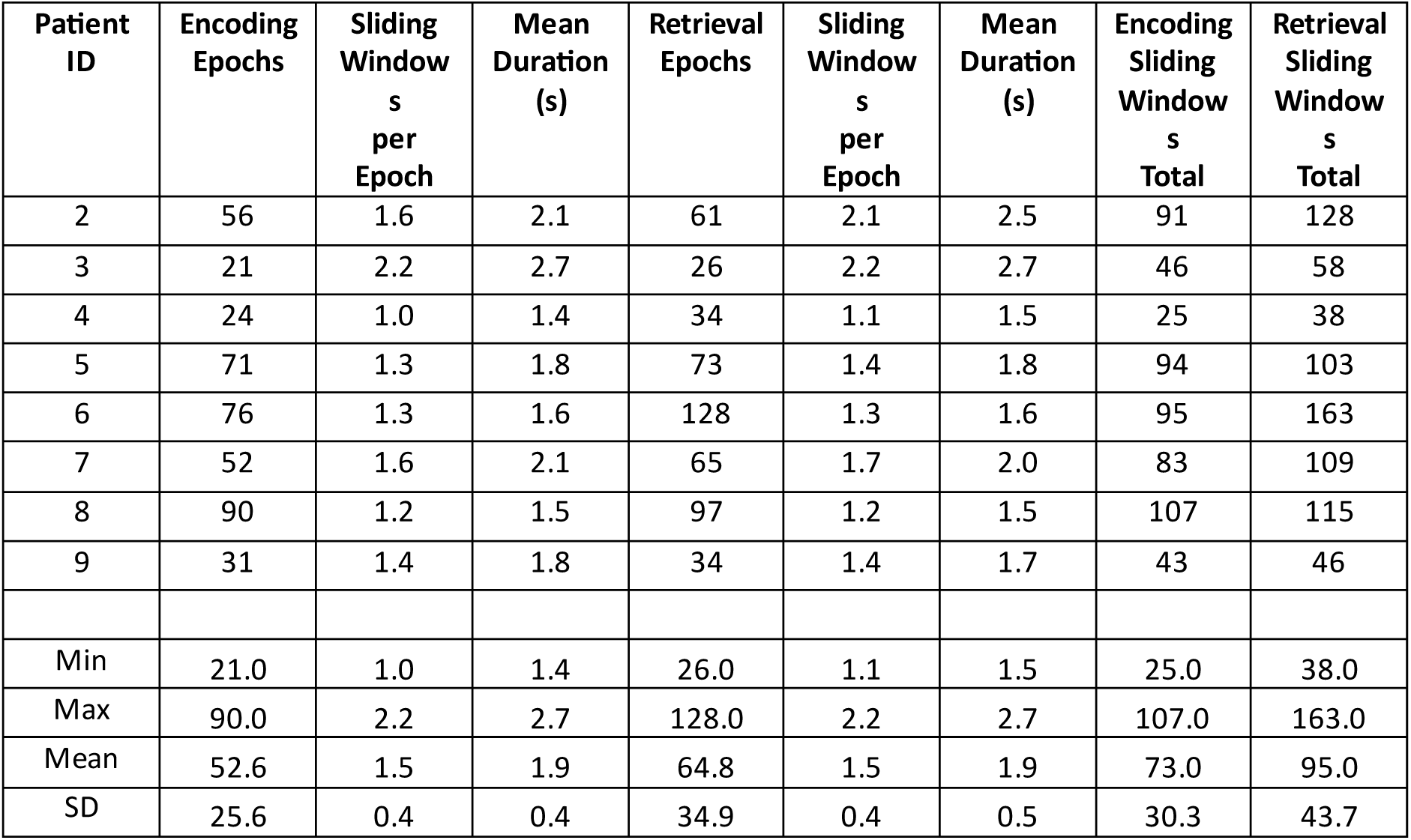
Task epochs for depth electrode contacts in the amygdala.

| Patient ID | Encoding Epochs | Sliding Windows per Epoch | Mean Duration (s) | Retrieval Epochs | Sliding Windows per Epoch | Mean Duration (s) | Encoding Sliding Windows Total | Retrieval Sliding Windows Total |
| --- | --- | --- | --- | --- | --- | --- | --- | --- |
| 2 | 56 | 1.6 | 2.1 | 61 | 2.1 | 2.5 | 91 | 128 |
| 3 | 21 | 2.2 | 2.7 | 26 | 2.2 | 2.7 | 46 | 58 |
| 4 | 24 | 1.0 | 1.4 | 34 | 1.1 | 1.5 | 25 | 38 |
| 5 | 71 | 1.3 | 1.8 | 73 | 1.4 | 1.8 | 94 | 103 |
| 6 | 76 | 1.3 | 1.6 | 128 | 1.3 | 1.6 | 95 | 163 |
| 7 | 52 | 1.6 | 2.1 | 65 | 1.7 | 2.0 | 83 | 109 |
| 8 | 90 | 1.2 | 1.5 | 97 | 1.2 | 1.5 | 107 | 115 |
| 9 | 31 | 1.4 | 1.8 | 34 | 1.4 | 1.7 | 43 | 46 |
| Min | 21.0 | 1.0 | 1.4 | 26.0 | 1.1 | 1.5 | 25.0 | 38.0 |
| Max | 90.0 | 2.2 | 2.7 | 128.0 | 2.2 | 2.7 | 107.0 | 163.0 |
| Mean | 52.6 | 1.5 | 1.9 | 64.8 | 1.5 | 1.9 | 73.0 | 95.0 |
| SD | 25.6 | 0.4 | 0.4 | 34.9 | 0.4 | 0.5 | 30.3 | 43.7 |

**Supplementary Table 4:**
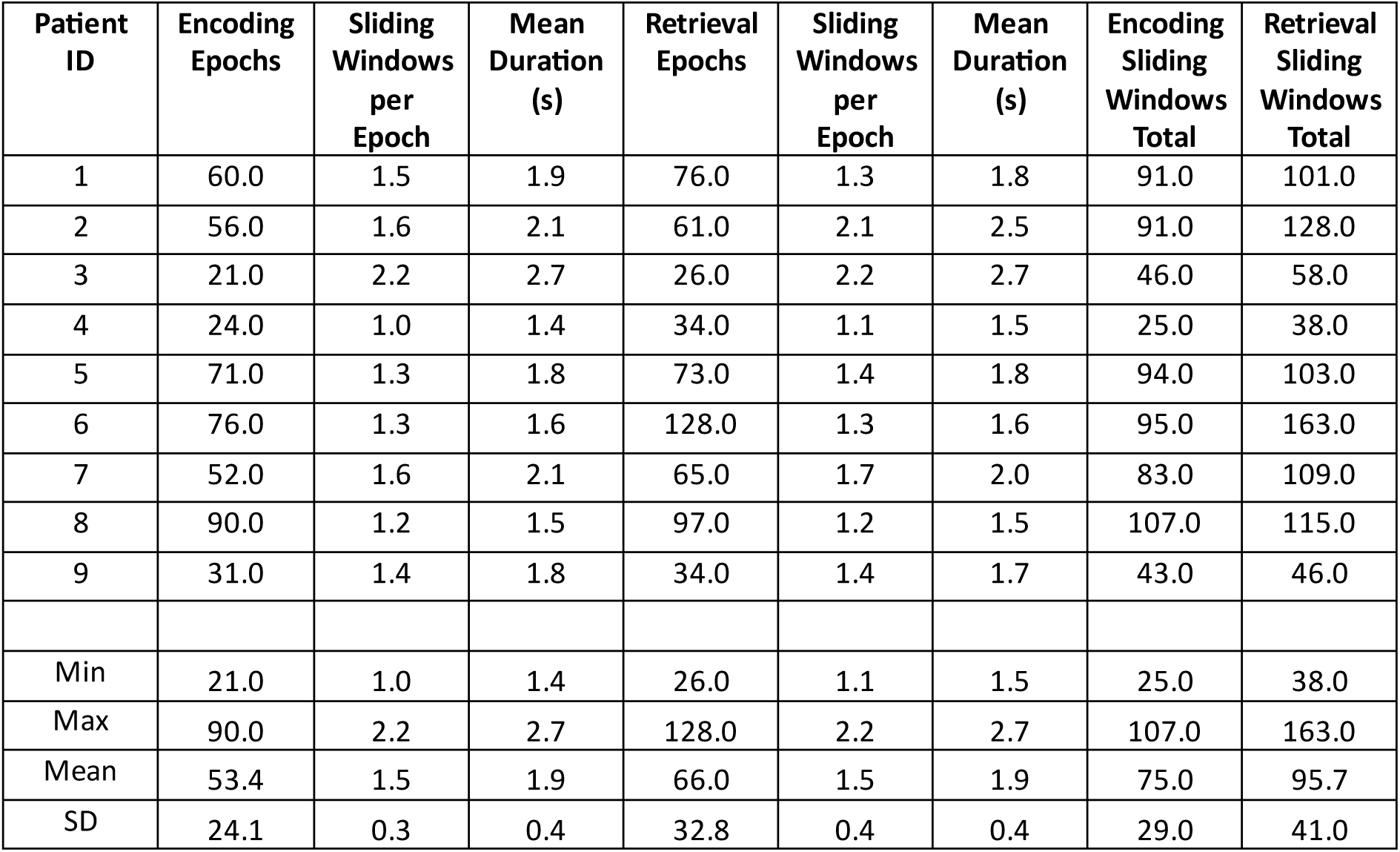
Task epochs for depth electrode contacts in the temporal neocortex.

## Notes

### Competing Interest Statement

The authors have declared no competing interest.

